# Mobility as the purpose of postural control

**DOI:** 10.1101/110932

**Authors:** Charlotte Le Mouel, Romain Brette

## Abstract

Counteracting the destabilizing force of gravity is usually considered to be the main purpose of postural control. However, from the consideration of the mechanical requirements for movement, we argue that posture is adjusted in view of providing impetus for movement. Thus, we show that the posture that is usually adopted in quiet standing in fact allows torque for potential movement. Moreover, when performing a movement - either voluntarily or in response to an external perturbation - we show that the postural adjustments are organized both spatially and temporally so as to provide the required torque for the movement. Thus, when movement is performed skillfully, the force of gravity is not counteracted but actually used to provide impetus to movement. This ability to move one’s weight so as to exploit the torque of gravity seems to be dependent on development and skill learning, and is impaired in aging.

## 1 Introduction

The position of the center of mass (CoM) is adjusted by the central nervous system during quiet standing (Sasagawa et al., 2009; Winter et al., 1998), in reaction to perturbations (Horak and Nashner, 1986), and in voluntary movement (Cordo and Nashner, 1982; Lee et al., 1990; Pedotti et al., 1989). The traditional theory is that the purpose of this postural control is to immobilize the center of mass despite movement and external perturbations (Bouisset and Do, 2008; Horak, 2006; Massion et al., 2004; Nashner et al., 1989). We will refer to this theory as the immobility theory. The underlying assumption is that, because of gravity, standing is unstable. Therefore, if the CoM is displaced from its equilibrium position, then the displacement must be counteracted by postural adjustments, so as to return the CoM to its equilibrium position, otherwise the person will inevitably fall. As argued by Hasan (2005), this notion stems from an analysis of how linear systems respond to perturbations: in linear systems, if deviations from the unique equilibrium position are not corrected, then they grow exponentially. Balance (the ability to prevent falling), is therefore assumed to be equivalent to stabilization, in the strict sense of immobilizing the CoM at a unique equilibrium position by counteracting any displacement away from this position. From this assumption, it follows that moving poses a threat to balance, since any voluntary movement might displace the CoM. This theory has motivated a large body of experiments, performed over the last thirty years, in which a subject is asked to perform a movement of the upper body, while their muscle activity is being recorded (Cordo and Nashner, 1982; Crenna et al., 1987; Lee et al., 1990; Pedotti et al., 1989). In these experiments, a change in the contraction of the lower leg muscles is systematically observed, and this change often precedes the contraction of the upper body muscles. This is interpreted by saying that movement of the upper body might displace the CoM, and must therefore be counteracted by the contraction of the lower leg muscles so as to immobilize the CoM despite movement.

We will argue however that the equivalence between balance and immobilization does not hold for human postural control, and that these postural responses should be understood as providing the impetus for the movement. We will indeed show that during quiet standing, voluntary movement, and in reaction to perturbations, the position of the CoM is not immobilized at a unique equilibrium position, but on the contrary adjusted so as to use the torque of one’s own weight, either to counteract external forces so as to maintain balance, or to provide impetus for voluntary movement. We therefore develop an alternative to the immobility theory. We propose that the purpose of postural control is mobility, the ability to produce appropriate impetus by adjusting the position of the CoM. We will refer to this theory as the mobility theory.

We will first show that the posture which is typically adopted in quiet standing allows for one’s weight to be used to provide impetus to potential movement, and that when the direction of the movement to be performed can be anticipated, the position of the CoM during stance is shifted in that direction. Secondly, we will show that, during voluntary movement, postural adjustments which are traditionally thought of as immobilizing the CoM despite movement should on the contrary be interpreted as displacing the CoM at the initiation of the movement, so that one’s own weight can be used to provide impetus to the movement. Finally, we will show that this ability to use displace one’s weight, rather than immobilize it, plays a crucial role when balance is upset by external forces.

## 2 Adjustment of posture during stance

### 2.1 The standing posture allows for mobility

#### 2.1.1 The standing posture requires tonic muscular contraction

When someone is asked to stand quietly, without further instructions, they typically maintain their CoM vertically aligned with the middle of the foot, a few centimeters forwards of the ankle joint (Schieppati et al., 1994). However, when requested to do so, a young, healthy person can maintain their CoM at positions up to 40 % of their foot length forwards of its typical position, and up to 20 % backwards (Schieppati et al., 1994). There is therefore no unique equilibrium position for the CoM in quiet standing, since a young, healthy person can maintain a range of standing postures without this posing a threat to balance.

If the position of the CoM were controlled only in view of counteracting the torque of one’s weight, then it would be most appropriate to place it vertically above the ankles, such that weight would exert no torque (Fig. 1A). This position can indeed be maintained with minimum lower leg muscle contraction (Schieppati et al., 1994). However, when no instructions are given, subjects maintain their CoM vertically aligned with the middle of the foot, a few centimeters forwards of the ankle joint (Fig. 1B), so that the weight exerts a forwards torque. In order to maintain this posture, an equivalent backwards torque must be exerted by the ground reaction force (see Appendix 6.1.1). As developed in the Appendix (6.1.2), the torque of the ground reaction force is determined by the contraction of the lower leg muscles. Indeed, if we consider the forces acting on the foot, the weight of the body, carried by the skeleton, is applied at the ankle and therefore exerts no torque. The ground prevents the foot from turning, therefore the ground reaction torque instantly opposes the torque exerted by the lower leg muscles onto the foot (Fig. 2). Maintaining a standing posture with the CoM forwards of the ankles therefore requires tonic contraction of the calf muscles. (Fig. 1B, Schieppati et al., 1994). The normal standing posture is therefore not the most economical in terms of muscular contraction.

**Fig. 1.**
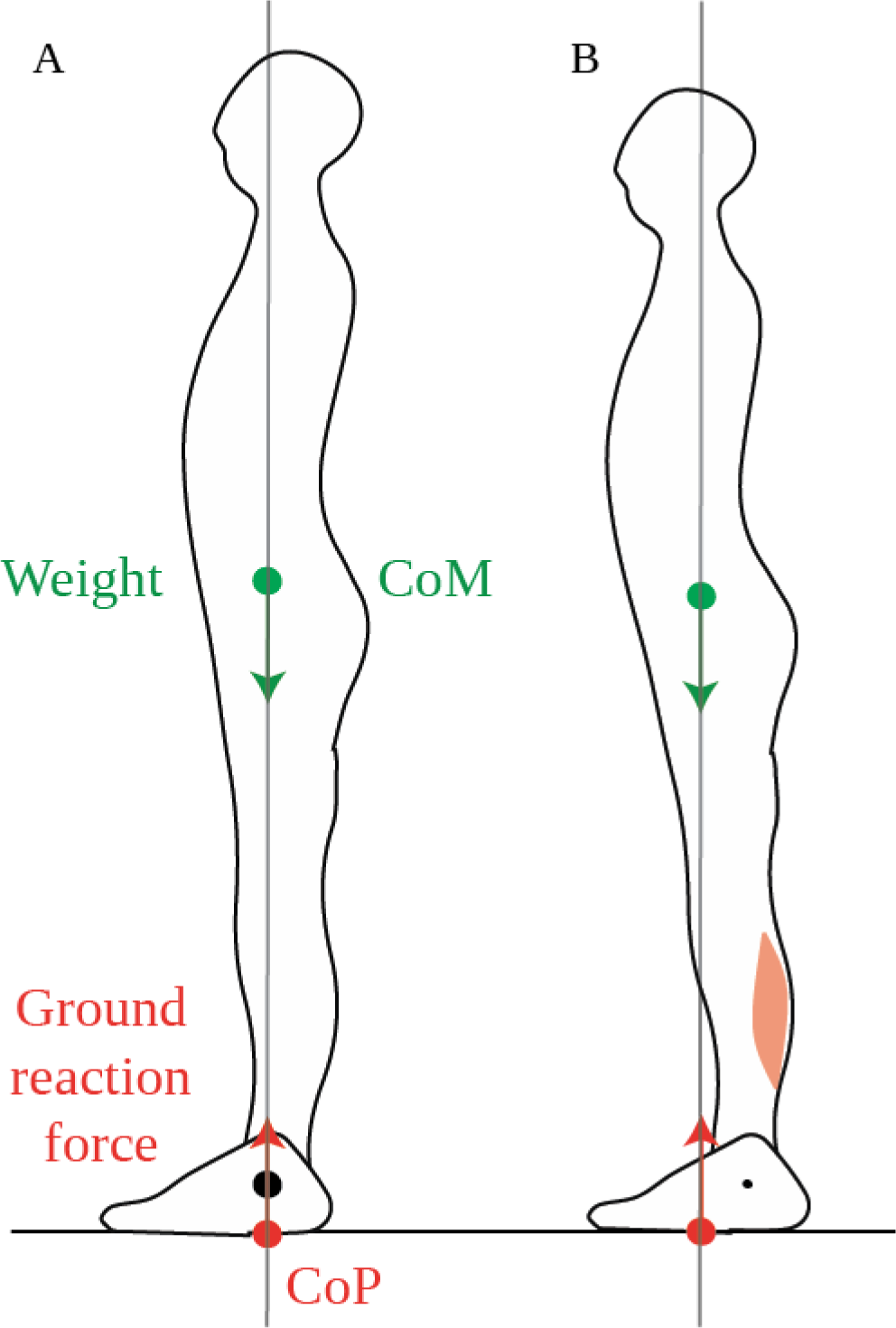
Standing posture. A. When the CoM (green dot) is vertically aligned with the ankle joint (black dot), the weight (green arrow) exerts no torque around the ankle. In order to maintain this posture, the ground reaction force (red arrow) must also exert no net torque around the ankle, therefore its point of application, the CoP (red dot) must also be vertically aligned with the ankle. B. In the typical quiet standing posture, the CoM is maintained forwards of the ankles, therefore weight exerts forwards torque around the ankles. This is compensated for by backwards torque of the ground reaction force, which requires tonic calf muscle contraction.

#### 2.1.2 The standing posture allows torque for movement

Why would subjects actively maintain their CoM forwards of the ankles in quiet standing if this is not efficient? We suggest that this allows them to use their own weight for initiating forwards movements. Forwards torque for movement can only be induced by the external forces: the person’s weight and the ground reaction force. As we have shown (Appendix 6.1.2) the ground reaction torque instantly follows the torque of the lower leg muscles. However, this torque is limited. Indeed, as long as the person neither jumps up nor collapses, the ground reaction force has the same magnitude as the person’s weight. Its torque is therefore the product of the weight, and the distance between the ankle and the point of application of the ground reaction force, called the center of pressure and noted CoP. Thus, contracting the calf muscles (gastrocnemius and soleus) shifts the CoP forwards of the ankle (Fig 2A), and contracting the shin muscle (tibialis anterior) shifts the CoP backwards of the ankle (Fig. 2C), but the CoP can only move within the limited range of the foot (see Appendix 6.1.3 for further detail).

**Fig. 2.**
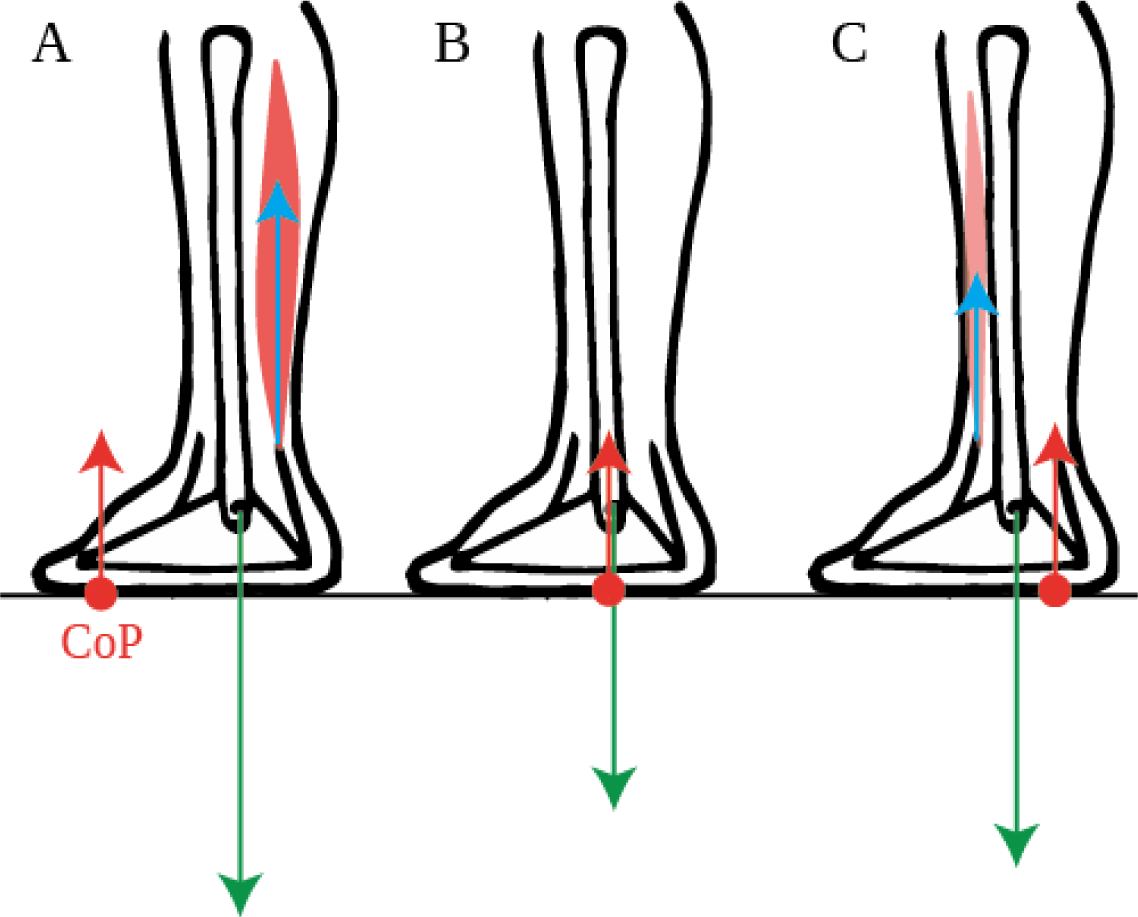
Torques exerted on the foot. The force exerted by the lower leg bones onto the foot (green arrow) exerts no torque around the ankle. The torque of the ground reaction force (red arrow) and of the forces exerted by the lower leg muscles onto the foot (blue arrow) are therefore opposite when the foot remains immobile: A. the torque around the ankles exerted by the calf muscles onto the foot is instantly compensated for by a forwards shift of the CoP (red dot). B. When the lower leg muscles exert no torque onto the foot, then the CoP is below the ankle C. The torque around the ankles exerted by the shin muscle onto the foot is instantly compensated for by a backwards shift of the CoP.

The net torque is proportional to the distance between the CoM and the CoP. Whereas the CoP moves instantly when the forces exerted by the muscles change, but can only move within the limited range of the foot, the position of the CoM on the other hand, does not change instantly when the forces exerted by the muscles change. This first requires the sum of the external forces to accelerate the CoM. Displacements of the CoM therefore occur more slowly than displacements of the CoP (as seen for example in Burleigh et al., 1994). Thus, the initial net torque that can be produced, either for opposing external perturbation forces or for voluntary movement, is limited by the initial position of the CoM (see Appendix 6.1.4 for further detail).

When initiating fast forwards movements, either starting to walk (Burleigh et al., 1994) or movements performed with the feet in place such as leaning forwards (Crenna et al., 1987) or rising onto one’s toes (Nardone and Schieppati, 1988), the CoP is first brought towards the heel by inhibiting the calf muscle contraction and contracting the shin muscle (Burleigh et al., 1994; Crenna et al., 1987; Nardone and Schieppati, 1988). If the CoM were initially above the ankle, this would produce little initial forwards torque (Fig. 3A), whereas with the CoM forwards of the ankle this produces larger torque (Fig. 3B). Maintaining the CoM forwards of the ankle thus allows one’s own weight to be used for initiating forwards movement. Maintaining the CoM in the middle of the foot allows for either forwards or backwards initial torque to be induced by changes in the forces of the lower leg muscles.

### 2.2 The standing posture is actively maintained

This position of the CoM is precisely and actively maintained on a short timescale, with small adjustments of the CoP in quiet standing serving to immobilize the CoM at this position (Fig. 4A, Winter et al., 1998). Moreover, the tonic contraction of the calf muscles is adjusted when standing on different slopes so as to maintain the CoM aligned with the middle of the foot (Fig. 4B, Sasagawa et al., 2009). This precise positioning is also maintained at the longer timescales of growth and aging. Indeed, the curvature of the spine and trunk increases with aging (red line in Fig. 4C, Schwab et al., 2006), and the position of the CoM is maintained across people with different trunk curvatures by shifting the position of the pelvis relative to the heels (Fig. 4C, Lafage et al., 2008; Schwab et al., 2006).

**Fig. 3.**
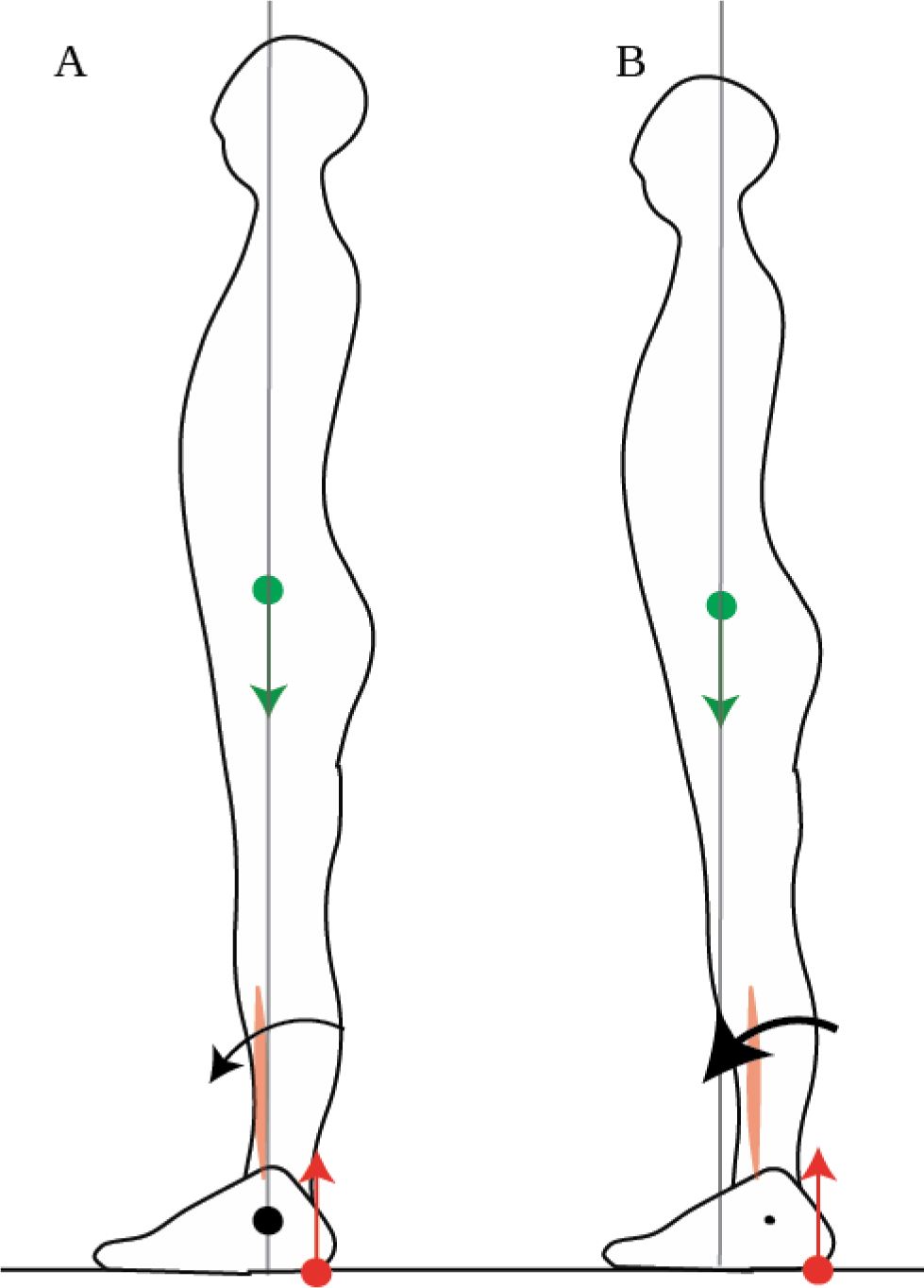
Net torque is limited by the position of the CoM. In order to initiate a forwards movement, the CoP is brought to the heel by inhibiting calf muscle contraction and contracting the shin muscle. When the CoM is vertically aligned with the ankle (A), the net forwards torque is small. When the CoM is forwards of the ankle (B), the net forwards torque is larger.

Moreover, this forwards position of the CoM emerges with skill learning. Thus, Clément and Rézette (1985) observed acrobats at various competitive levels performing handstands. All the acrobats were able to maintain their balance in the upside-down posture, however they did so in different ways. The acrobats at lower competitive levels maintained their mean CoP a few millimetres forwards of their wrist; they could therefore maintain their posture with very little tonic contraction in the arm muscles (Fig. 4D, left). The acrobats at higher competitive levels maintained their mean CoP more forwards of their wrists, with the acrobat at the highest level maintaining his mean CoP 3 cm forwards of his wrists; this posture requires tonic contraction of the wrist extensors (Fig. 4D, right).

Thus, the standing posture is actively adjusted so as maintain the CoM above the middle of the foot (and above the middle of the hand in handstands). Contrary to the immobility theory, this position is not a unique equilibrium point, since a variety of standing postures can be maintained without this leading to a loss of balance. According to the mobility theory, this position is maintained because it allows for torque of the appropriate direction to be produced at short notice, even when this direction cannot be anticipated. This may be useful both for opposing external perturbations and for initiating voluntary movements.

**Fig. 4.**
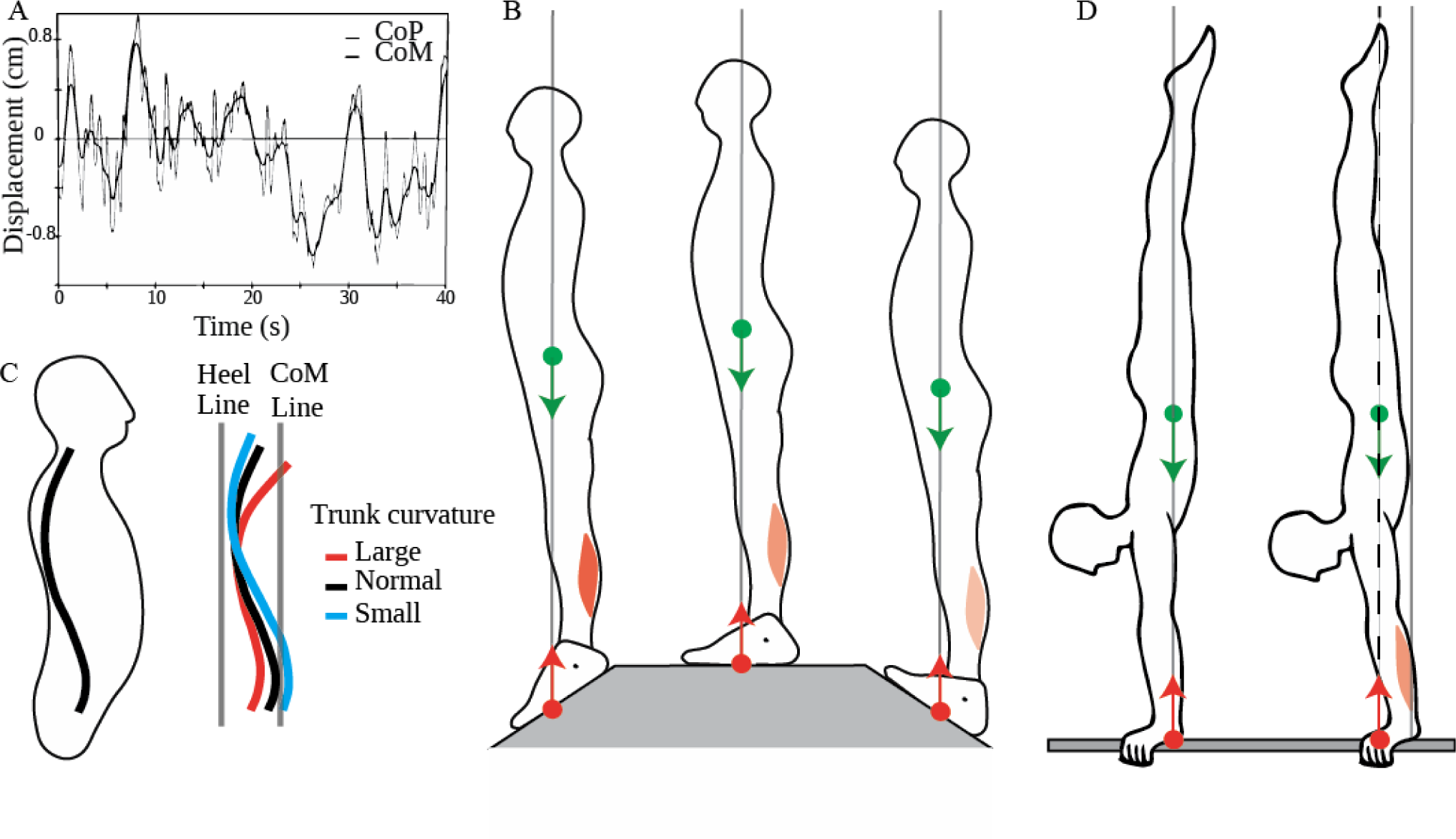
Adaptation of the position of the CoM. A. Displacement of the CoP and CoM in quiet standing as a function of time, adapted from Winter et al. (1998): small ongoing shifts of the CoM are tracked and overtaken by shifts in the CoP. B. The tonic calf muscle contraction decreases when going from a slope with the toes down (left panel), to a flat slope (middle panel), to a slope with the toes up (right panel) such that position of the CoM is maintained vertically aligned with the middle of the feet. C. People of different trunk curvatures maintain their pelvis at different distances from the heel line (vertical line above the heel), such that the CoM line (vertical line passing through the CoM) is at the same distance from the heel line. D. Left panel: Acrobats at lower competitive levels maintain their CoP and CoM aligned with their wrist without tonic contraction of their wrist extensors. Right panel: Acrobats at higher competitive levels maintain their CoP and CoM forwards of their wrist, through tonic contraction of their wrist extensors.

### 2.3 The standing posture is adjusted in anticipation of movement

When the direction of the appropriate torque can be anticipated, the mobility theory predicts that the CoM would be displaced in that direction in anticipation of the movement. Such a shift can indeed be induced experimentally, either by challenging someone’s balance in a predictable direction, or by indicating in advance the direction of a voluntary movement to be performed.

**Fig. 5.**
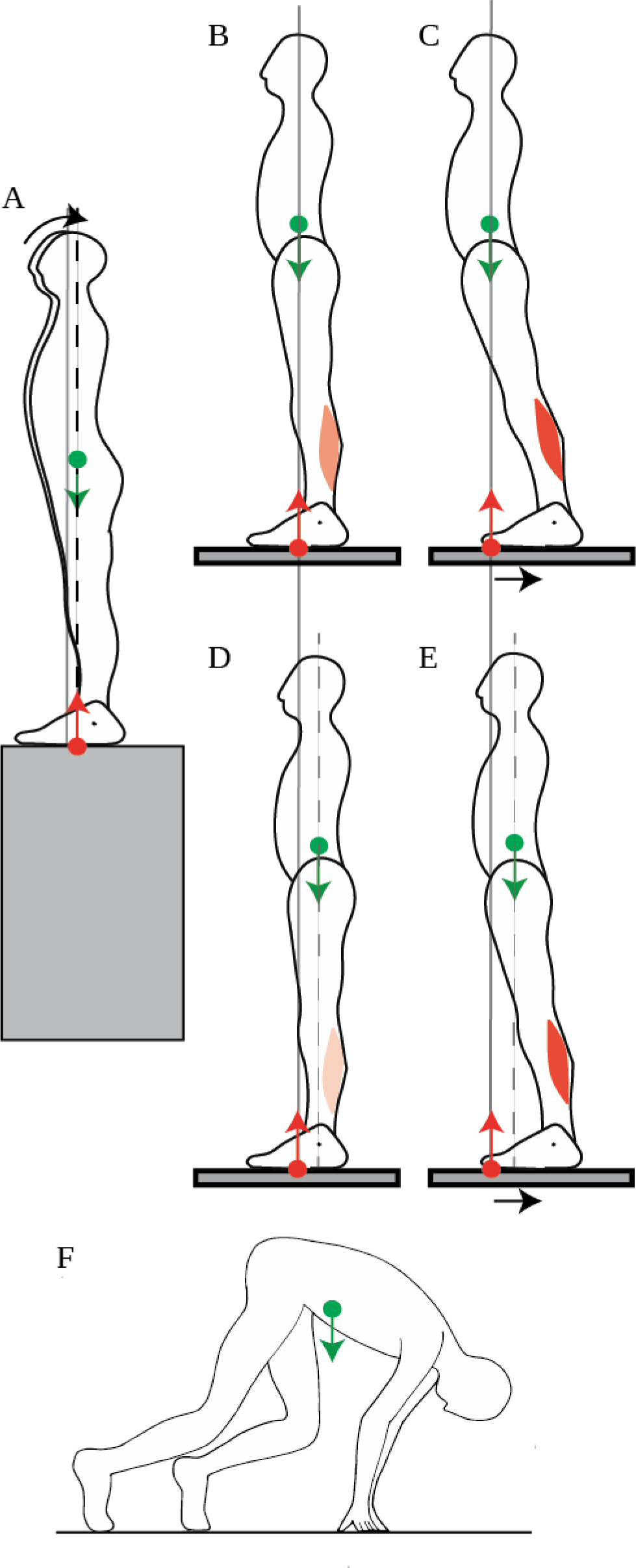
Adjustment of the position of the CoM. A. When a person stands facing a slope, they shift their CoM slightly backwards. B-C: When someone stands normally (B) and the platform they stand on is shifted backwards, their CoM ends up far forwards of the ankle joints, which limits the net backwards torque for straightening up (C). D-E: When a backwards perturbation is repeated, the person shifts their CoM backwards in quiet standing (D), which increases the net backwards torque for straightening up after the perturbation (E). F: In the posture adopted before a sprint, the CoM is placed far forwards of the feet by having the arms carry some of the weight.

Someone’s balance can be challenged by having them stand facing the edge of the platform they are on. According to the immobility theory, this should lead, if anything, to an even more stringent immobilization of the CoM at its equilibrium position, but what is observed is that the CoM is shifted slightly backwards (Fig. 5A, Carpenter et al., 2001). This is in accordance with the mobility theory, since it increases the person’s capacity for producing backwards torque, in the eventuality that they might be subjected to a forwards push. In the experiment, the person’s balance was not challenged beyond placing them in front of a drop, which might explain why the shift in CoM position was rather small (less than a centimeter).

Another way of challenging someone’s balance is to have them stand on a platform (Fig. 5B) which is then translated backwards (Fig. 5C). The person ends up with their CoM in a forward position relative to the feet. A commonly observed response to such a translation is to straighten up (Welch and Ting, 2014). This requires backwards torque, however their capacity for producing backwards torque is limited by the forwards position of their CoM (Fig. 5C). If such a perturbation is repeated, then over a few trials, the person adjusts their quiet standing posture by shifting their CoM backwards by a few centimeters (Fig. 5D, Welch and Ting, 2014). This is again in contradiction with the immobility theory, but in accordance with the mobility theory, since the backwards shift of the CoM increases the person’s capacity to produce backwards torque for straightening up (Fig. 5E). When the platform is repeatedly translated forwards, then the person shifts their CoM forwards (Welch and Ting, 2014).

The mobility theory predicts that the position of the CoM in quiet standing would also be shifted if the direction in which a voluntary movement to be performed could be anticipated. This occurs at the start of a race: in sprinting, the initial forwards acceleration is crucial in winning the race. Consistently with the mobility theory, the CoM in the starting position is shifted even beyond the toes by several tens of centimeters (Slawinski et al., 2010). This is achieved by placing the hands on the ground and having the hands carry some of the weight (Fig. 5F). This ability to use one’s own weight to produce torque for movement again seems to depend on skill learning. Indeed, in elite sprinters, the CoM is shifted 5 centimeters further forwards than for well-trained sprinters (Slawinski et al., 2010).

### 2.4 Summary

Thus, when the direction of the appropriate torque to be produced cannot be anticipated, the CoM is positioned at the middle of the feet, in a position which allows for both forwards and backwards torque to be produced. When the direction of the torque to be produced can be anticipated, then the standing posture is adjusted by shifting the CoM in that direction. This adaptation of the standing posture in view of movement seems to be dependent on learning.

## 3 Adjustment of posture during voluntary movement

According to the immobility theory, when a voluntary movement is being performed, postural control serves to immobilize the CoM despite the movement or the perturbation. The mobility theory predicts, on the contrary, that the position of the CoM is adjusted so as to use the torque of weight for movement. It therefore predicts that muscular contractions are temporally organized so as to accelerate the CoM at the initiation of the movement in the appropriate direction for producing torque for movement.

**Fig. 6.**
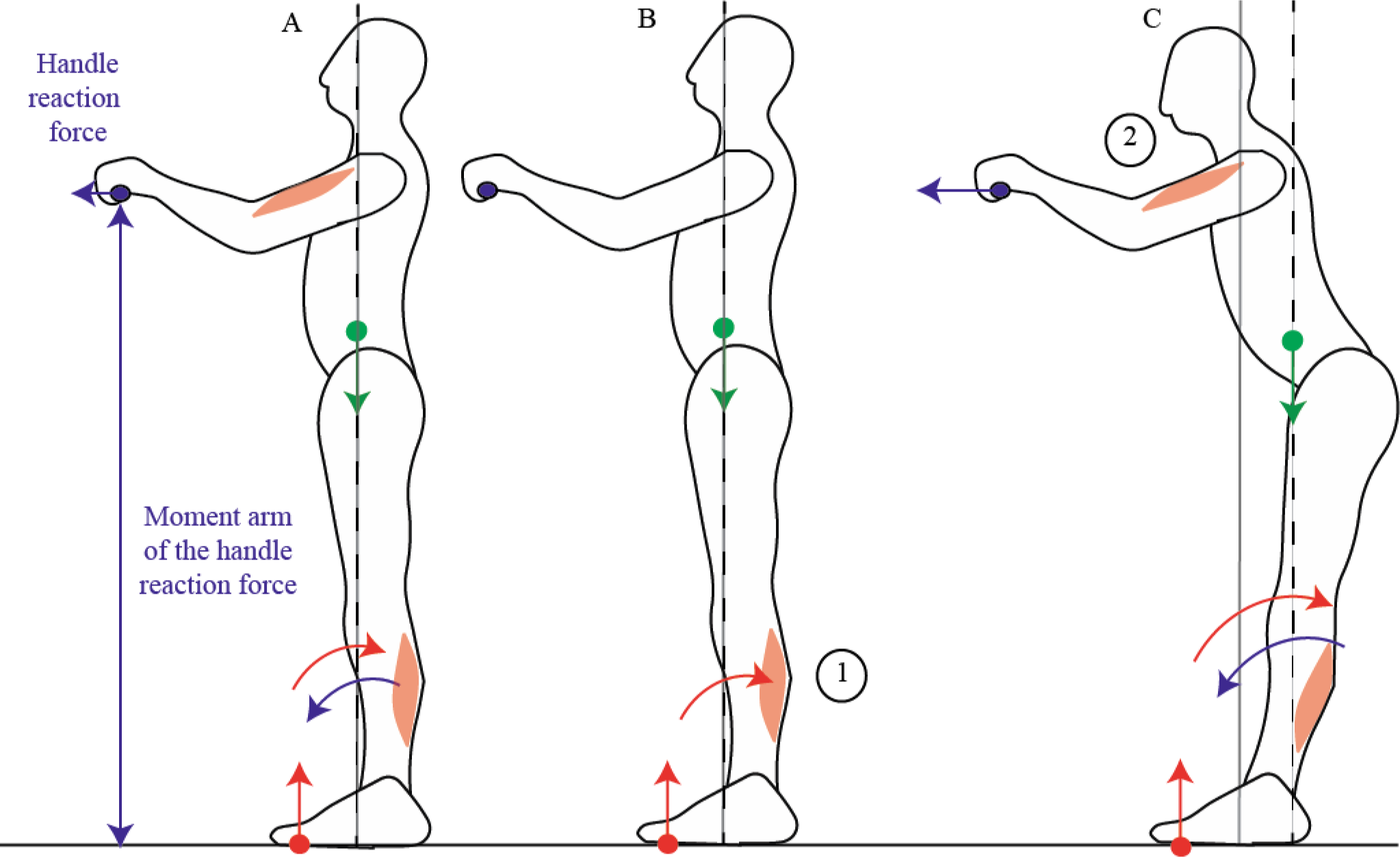
Pulling on a handle. When pulling on a handle, the handle reaction force (blue arrow) exerts forwards torque around the ankles which can compensated for by contracting the calf muscles (A). In preparation for pulling on a handle, subjects contract their calf muscles before their arm muscles (B), which displaces their CoM backwards, allowing for a larger net backwards torque to be exerted during the handle pull (C).

### 3.1 Initiation of voluntary movement

#### 3.1.1 Pulling on a handle

When someone pulls on a handle placed in front of them, the contraction of the arm muscles is preceded then accompanied by the contraction of the calf muscles (Cordo and Nashner, 1982; Lee et al., 1990). Cordo and Nashner (1982) suggest that this contraction of the calf muscles allows for the CoM to be immobilised despite the movement. However, in order for the CoM to be immobilised, the ground reaction torque would have to exactly compensate for the handle reaction torque throughout the movement, and this would notably require the calf and arm muscle contractions to be simultaneous (as in Fig. 6A). On the contrary, the initial contraction of the calf muscles which is observed (Cordo and Nashner, 1982) accelerates the CoM backwards (Fig. 6B, C, further details are provided in Appendix 6.2); and when the person is asked to pull harder on the handle, this initial period lasts longer, the calf muscle activation is stronger, and the initial backwards acceleration of the CoM is larger (Lee et al., 1990). This is in accordance with the mobility theory, since initially accelerating the CoM backwards allows one’s own weight to be used to assist the movement (Fig. 6C).

**Fig. 7.**
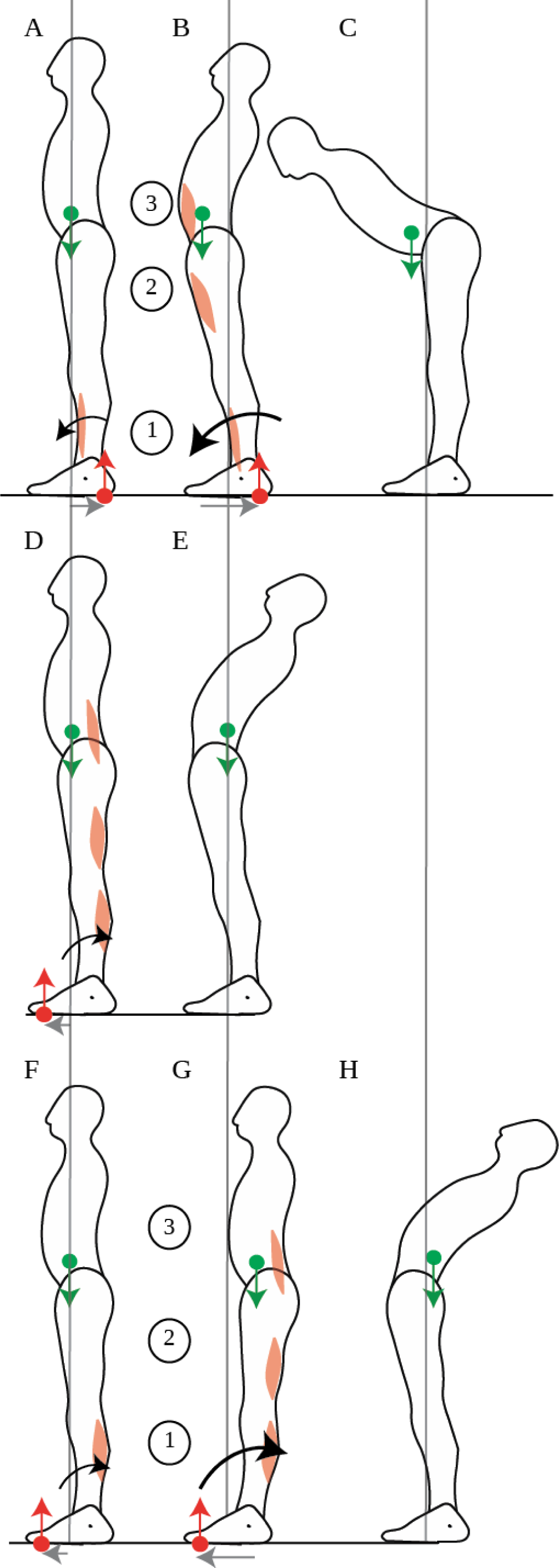
Leaning the trunk. When control subjects perform fast forwards leaning, the initial contraction of the shin muscle (A) accelerates the CoM forwards, thus allowing for more net forwards torque during the subsequent contraction of the ventral muscles (B), which enables the person to lean the trunk (C). When control subjects perform fast backwards leaning, the dorsal muscles contract simultaneously (D), which increases backwards rotational momentum without translating the CoM (E). When gymnasts perform fast backwards leaning, the initial contraction of the calf muscles (F) accelerates the CoM backwards, thus allowing for more net backwards torque during the subsequent contraction of the dorsal muscles (G) which enables the gymnast to lean the trunk (H). The sequence of activation of the muscles is indicated by the numbers 1 to 3.

#### 3.1.2 Leaning the trunk

When someone leans the trunk forwards, the contraction of the abdominal muscles is preceded then accompanied by the inhibition of calf muscle contraction and the contraction of the shin muscle (Fig. 7A-C; Crenna et al., 1987). The CoM could in theory be immobilized if the shin and abdominal muscle contractions were simultaneous, such that the forwards acceleration of the CoM induced by the shin muscle contraction would compensate for the backwards acceleration of the CoM induced by the abdominals contraction (further details are provided in Appendix 6.2), as suggested by Alexandrov and colleagues (2001). However, these authors report an initial backwards displacement of the CoP (Fig. 7A), followed by a forwards displacement of the CoM (Fig. 7B), in accordance with the sequential muscular contraction observed by Crenna and colleagues (1987). This contradicts the immobility theory, but concords with the mobility theory’s predictions.

Thus, postural responses should be considered as an integral part of the movement itself, since they provide the torque for the movement, first by shifting the CoP and secondly by accelerating the CoM through sequential muscle contraction (a more complete explanation can be found in Appendix 6.2).

#### 3.1.3 Gait initiation

Bouisset and Do (2008) distinguish between two types of anticipatory postural adjustments. For voluntary movements without a change in the basis of support, such as raising the arm, they provide a very classical interpretion for the displacement of the CoM which precedes the displacement of the arm. They present it as a counterperturbation whose purpose is to “counterbalance the disturbance to postural equilibrium due to the intentional forthcoming movement” (Bouisset and Zattara, 1981). However, for voluntary movements involving a change of the basis of support, such as walking, or rising onto one’s toes, they present anticipatory postural adjustments as a perturbation involved in “body weight transfer” (Do et al., 1991).

We propose that in movements with or without a change in the basis of support, anticipatory postural adjustments play the same role of moving the CoM in order to provide impetus for movement. Indeed, the changes in posture which precede walking are organised in the same way as those which precede pulling on a handle or leaning the trunk. Thus, when going from standing to walking, a few hundred milliseconds before the heel of the swing foot is raised, the calf muscles are silenced and the shin muscle contracts, which brings the CoP to the heels and accelerates the CoM forwards, even before the first step is taken (Fig 1D; Burleigh et al., 1994). This is in accordance with the mobility theory, since initially accelerating the CoM forwards allows one’s own weight to be used to assist the movement. Indeed, this initial acceleration of the CoM is correlated with the speed reached at the end of the first step, and is larger if the person is asked to walk faster (Brenière et al., 1987).

### 3.2 The ability to use one’s weight for movement requires practice

For walking, a movement which is learned very early on in life, the ability to displace the CoM at the initiation of the movement emerges over the course of development (Bril et al., 2015; Ledebt et al., 1998). The amplitude of the initial backwards shift of the CoP thus increases over the first several years of life as children learn to walk faster (Bril et al., 2015; Ledebt et al., 1998). It then decreases with age, and with certain neurological diseases such as Parkinson’s disease (Halliday et al., 1998; Mancini et al., 2016).

**Fig. 8.**
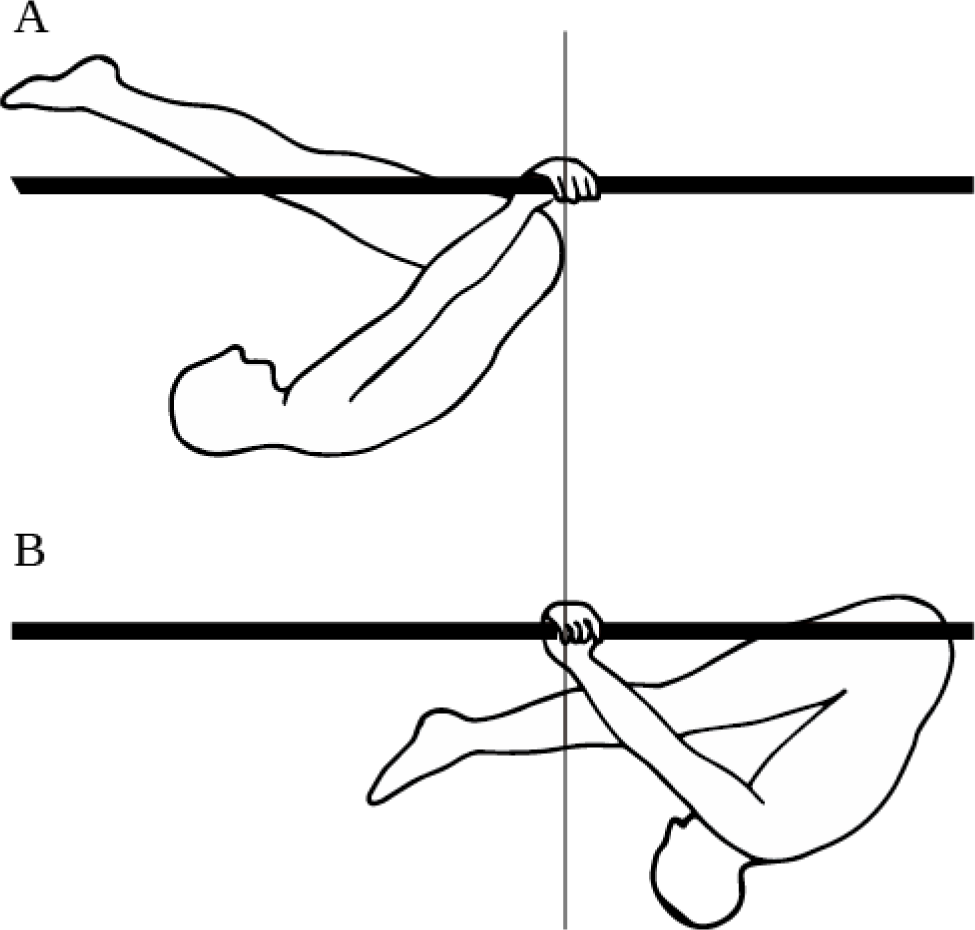
Gymnastics skill: swings under parallel bars. A. Forwardmost position in the swing. B. Backwardmost position in the swing.

For leaning the trunk, the sequential muscle contraction, which allows for the displacement of the CoM at the initiation of the movement, seems to be dependent on learning. Indeed, when control subjects are asked to lean backwards, a movement for which they presumably have less practice than leaning forwards, then the calf and dorsal trunk muscle contractions are simultaneous (Fig. 7D), and the movement is performed twice as slowly as leaning forwards (Pedotti et al., 1989). However, when gymnasts are asked to lean backwards, then their calf muscles contract first, and they perform the movement faster than controls (Fig. 7F-H, Pedotti et al., 1989). Moreover, the ability to displace one’s CoM during movement seems to remain plastic throughout life, and to depend on the possibility to use one’s weight to assist movement. Thus, when astronauts return from a several months journey in space (during which they could not use their weight to assist their movements), the forwards displacement of the CoM when leaning forwards is no longer observed (Baroni et al., 2001).

Finally, for movements requiring skill learning, the temporal coordination which enables using one’s weight to provide impetus for movement seems to develop with skill learning. Thus, when learning a complex gymnastics skill, such as the swings under parallel bars, in bent inverted hang position (Fig. 8), beginners swing their legs and arms in synchrony, whereas experts swing their legs out of phase with their arms, which allows them to use the work of their own weight to provide impetus to the swing (Delignières et al., 1998).

## 4 Balance requires mobility rather than immobility

According to the immobility theory, if postural control does not immobilize the CoM at a unique equilibrium position, then the person must fall (Bouisset and Do, 2008; Horak, 2006; Massion et al., 2004; Nashner et al., 1989). We have shown however that in quiet standing, people can keep their balance over a range of positions of the CoM (Schieppati et al., 1994), and actually displace their CoM when their balance is challenged in a predictable (Carpenter et al., 2001; Welch and Ting, 2014). Moreover, we have shown that in well-practiced movements, people accelerate their CoM at the initiation of the movement, without this leading to a loss of balance (Cordo and Nashner, 1982; Crenna et al., 1987; Lee et al., 1990; Pedotti et al., 1989). We will now show that the response to an external perturbation should be considered as a movement in its own right, and therefore also benefits from the ability to use one’s weight for movement, rather than to immobilize it.

**Fig. 9.**
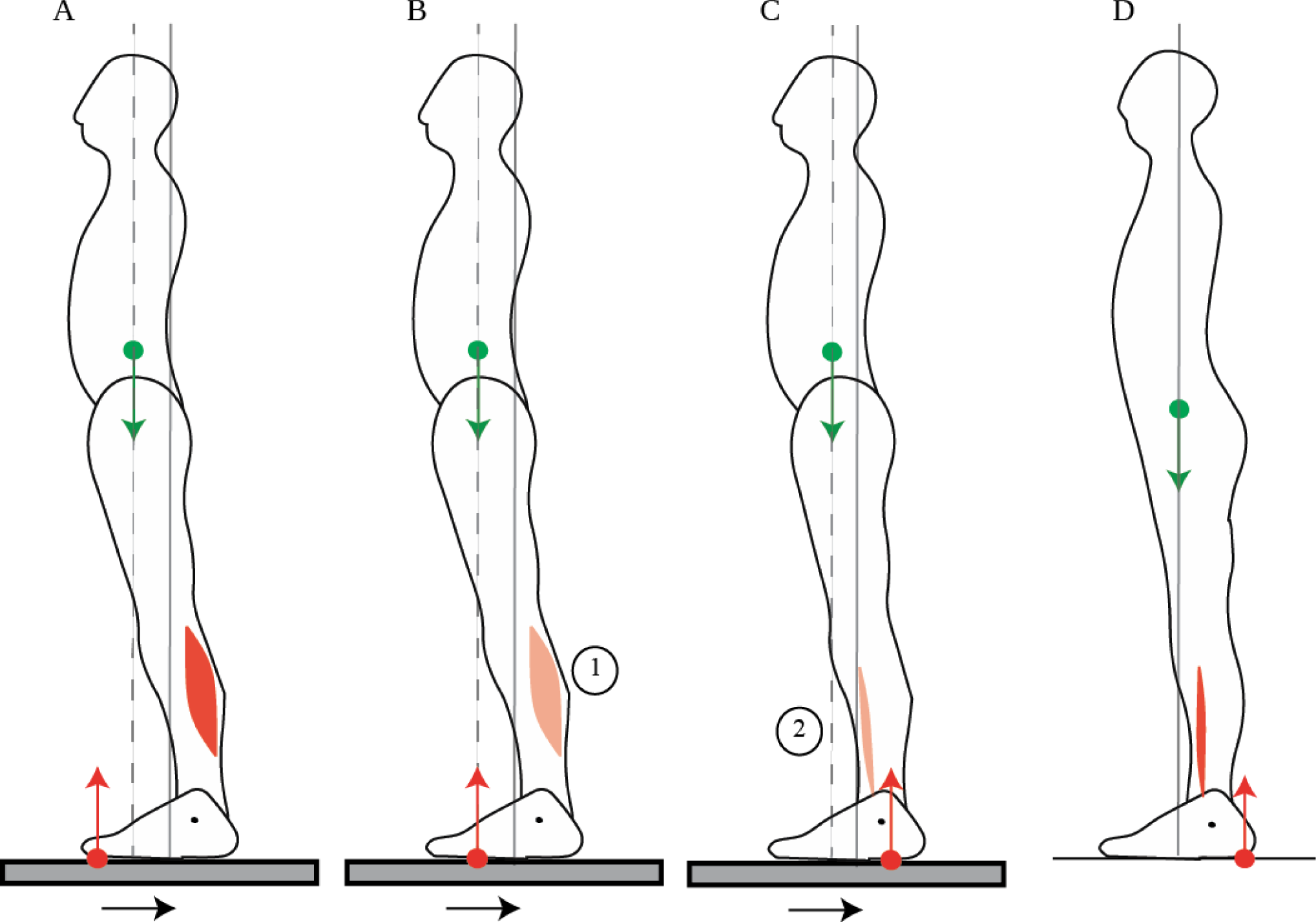
Response to platform translation. When straightening up after a platform translation (A), the initial contraction of the calf muscle accelerates the CoM backwards which increases the potential net backwards torque. When stepping forwards in response to a platform translation (B-C), this initial calf muscle contraction is reduced (B). Then, the shin muscle contracts (C). This shin muscle contraction is smaller than the initial contraction of the shin muscle which accelerates the CoM forwards when the person steps forwards without a platform translation (D). The sequence of activation of the muscles is indicated by the numbers 1 to 2.

### 4.1 Responding to external perturbations

#### 4.1.1 Straightening up after a platform translation

When the platform on which someone stands is translated backwards, the CoM ends up in a forward position relative to the feet (Fig. 9A,B), as seen in section 2.3. A response which is commonly observed is to straighten up (Horak and Nashner, 1986). The backwards acceleration of the CoM is performed through a sequential contraction of the dorsal muscles, starting with the calf muscles (Fig. 9A), then the dorsal thigh then dorsal trunk muscles (Horak and Nashner, 1986). This contraction pattern is usually not considered as an actual movement, since it moves the CoM closer to its initial position, in accordance with the immobility theory. However, we believe it should be considered as a movement in its own right. Indeed, straightening up after a platform translation requires producing the appropriate backwards torque. The sequential contraction pattern allows for the CoM to be initially accelerated backwards, which increases the net backwards torque for the movement. Further details are provided in Appendix 6.2. Moreover, contrary to the immobility theory, returning the CoM to its initial position is not the only way of preventing a fall.

#### 4.1.2 Stepping after a platform translation

Indeed, another response which is also commonly observed is to take a step forwards (Maki et al., 2003): the CoM is then not returned to its initial position, without this causing a loss of balance. This response takes advantage of the forwards position of the CoM, such that the CoM needs not be accelerated backwards, and indeed the initial calf muscle contraction and forwards CoP shift is much reduced (Fig. 9B) compared to when the person straightens up (Fig. 9A); nor does the CoM need to be accelerated forwards, and indeed the shin muscle contraction lasts much less long and the backwards shift of the CoP is much smaller (Fig. 9C; Burleigh et al., 1994) than when the person takes a step without the platform translation (Fig. 9D).

### 4.2 Emergence over development and impairment with aging

The ability to mobilize one’s weight emerges over development. Thus, when straightening up after a backwards platform translation, both the systematic recruitment of the dorsal muscles and their temporal sequencing emerge during development. They are not observed in prewalking infants, but are seen in children with a few years’ walking experience (Burtner et al., 1998).

This ability is then deteriorated with aging, and with Parkinson’s disease. The elderly, and even more so Parkinsonian patients, are less capable of moving their CoM, either when asked to adjust their quiet standing posture by leaning forwards or backwards (Schieppati et al., 1994), or during voluntary movement, such as gait initiation (Halliday et al., 1998). They are however quite as capable as young healthy adults of remaining immobile in quiet standing (Schieppati et al., 1994), and adjust the position of their pelvis to compensate for trunk curvature such that the CoM remains above the middle of the feet (Schwab et al., 2006). Nevertheless, they have a heightened risk of falling. Thus, although the elderly and Parkinsonian subjects are quite as capable as young adults of maintaining their CoM immobile during quiet standing, we suggest that their higher risk of falling is due to a limited capacity to move when this becomes necessary to prevent a fall. Therefore, not only is immobilizing the CoM unnecessary for balance, it moreover seems that balance benefits from the ability to move one’s CoM. This suggests that efficient balance training for the elderly can be achieved by practicing mobility (Xu et al., 2005).

## 5 Discussion

### 5.1 Posture is adjusted in view of mobility rather than immobility

Although the position of the CoM is adjusted by the nervous system, this postural control does not serve to immobilize the CoM. On the contrary, the position of the CoM is adjusted so as to use the torque of one’s own weight both for self-initiated movements and for responding to external perturbation forces.

Thus, in quiet standing, when the direction of the torque to be produced cannot be anticipated, the CoM is maintained above the middle of the foot (Schieppati et al., 1994), allowing for the torque of one’s weight to be used both for forwards and backwards movements. This position is actively maintained despite short-term changes in slope (Sasagawa et al., 2009) or long-term changes in trunk curvature (Schwab et al., 2006). However, when the direction of the torque to be produced can be anticipated, then the CoM is shifted in that direction. There is thus a small backwards shift of the CoM when someone is placed in front of a drop (Carpenter et al., 2001), or on a platform which is repeatedly translated backwards (Welch and Ting, 2014). Skill learning leads to much larger shifts in the position of the CoM, with the CoM placed forwards of the feet in anticipation of sprinting (Slawinski et al., 2010).

Moreover, during movement, we have shown that the postural responses which were thought to immobilize the CoM despite movement are actually temporally organized so as to accelerate the CoM at the initiation of the movement, in the appropriate direction such that the torque of one’s weight can be used for the movement (Cordo and Nashner, 1982; Crenna et al., 1987; Lee et al., 1990; Pedotti et al., 1989). These postural responses should therefore be understood as providing impetus to the movement.

Finally, we have shown that in order to respond effectively to external perturbation forces, the CoM need not be immobilized, since the person can take a step (Maki et al., 2003). When the person straightens up without taking a step (Horak and Nashner, 1986), this requires producing forces to counteract the external perturbation, and may benefit from the ability to mobilize one’s CoM rather than immobilize it. Balance therefore requires mobility rather than immobility.

### 5.2 Mobility emerges through development and skill learning

The ability to use one’s weight for movement emerges through development and skill learning, and remains plastic throughout life. The appropriate temporal organization of muscular contraction emerges during development both for walking and for balancing responses (Burtner et al., 1998; Ledebt et al., 1998). It is not observed for less practiced movements, such as when control subjects lean the trunk backwards (Crenna et al., 1987). The extent to which the CoM can be mobilized seems to depend on the level of skill: thus, both for sprinters at the initiation of a race (Slawinski et al., 2010) and acrobats performing handstands (Clément and Rézette, 1985), elite athletes place their CoM further forwards than well-trained athletes. Future work should address the following questions: how is this ability learned through development and practice? Does the impairment of this ability in aging result from a lack of practice, and could this ability be maintained during aging through appropriate training regimes?

## 6 Appendix

### 6.1 Torque induced by muscular contraction

#### 6.1.1 Torques of the external forces

When someone is standing on the ground, there are two external forces exerted on them: the person’s weight and the ground reaction force.

Gravity exerts a downwards vertical force whose magnitude is the person’s weight (their mass times the gravity of Earth). The point of application of the person’s weight is called the centre of mass and noted CoM. If the CoM is vertically aligned with the ankles, then the person’s weight does not exert a torque around the ankles (Fig. 1A). If it is forward of the ankles, then the weight exerts a forwards torque which is equal to the weight times the horizontal distance between the CoM and the ankles (Fig. 1B).

The ground supports the person’s weight, therefore, as long as the CoM remains at the same height, the vertical component of the ground reaction force is of equal magnitude but of opposite direction to the person’s weight. The ground reaction torque is therefore equal to the weight times the horizontal distance between the ankles and the point of application of the ground reaction force, called the centre of pressure and noted CoP.

The net torque around the ankles is thus determined by the horizontal distance between the CoP and the CoM: if they are vertically aligned, there is no net torque (Fig. 1 A and B), if the CoP is forwards of the CoM, then there is a net backwards torque (Fig. 1 C and D), and if the CoP is backwards of the CoM, then there is a net forwards torque around the ankles.

Such torque induces a change in the person’s rotational momentum around their ankles, which is the sum over their body segments of the segment’s mass, times its distance to the ankle, times its rotational speed (its speed perpendicularly to the axis joining it and the ankle).

#### 6.1.2 Lower leg muscle contraction changes the ground reaction torque

We will show that only the forces exerted by the lower leg muscles onto the foot may change the ground reaction torque around the ankles.

In order to understand how the internal forces induced by muscular contraction may affect the ground reaction force, we shall decompose the body and consider only the foot (Fig. 2). If the foot is on a rigid support and does not slip, then it can neither translate, nor rotate around the ankle. Therefore, both the sum of forces and the sum of torques around the ankle must be zero. The forces exerted onto the foot are the ground reaction force (see the red arrow in Fig. 2), the foot’s weight (which is negligible compared to the other forces), and the forces exerted by the lower leg onto the foot through on the one hand the muscles which attach onto the foot (see the blue arrows in Fig. 2), and on the other hand the bones, which exerts no torque around the ankles since it is applied at the ankles (see the green arrow in Fig. 2).

Thus, as long as the ground prevents the foot from moving, the torque of the ground reaction force around the ankle is exactly the opposite of that of the muscles of the lower leg. When the calf muscles contract, this pulls the heel upwards through the Achilles tendon (Fig. 2A). If the foot were in the air, it would rotate around the ankle joint bringing the toes down. Since the foot is against rigid ground, the ground resists the rotation of the foot by exerting backwards torque on the foot. Thus, any increase in the force that the calf muscles exert on the heel is instantly translated into an increase in the backwards torque of the ground reaction force on the entire body. As we have seen, as long as the CoM remains at the same height, the vertical component of the ground reaction force is of equal magnitude but of opposite direction to the person’s weight. Since the magnitude of the vertical component of the ground force does not change, contraction of the calf muscles can only induce backwards ground reaction torque by shifting the CoP forwards (Fig. 2A). Likewise, any increase in the force of the shin muscle is instantly translated into an increase in the forwards torque of the ground reaction force on the entire body, through a backwards shift in the CoP (Fig. 2C).

**Fig. 10.**
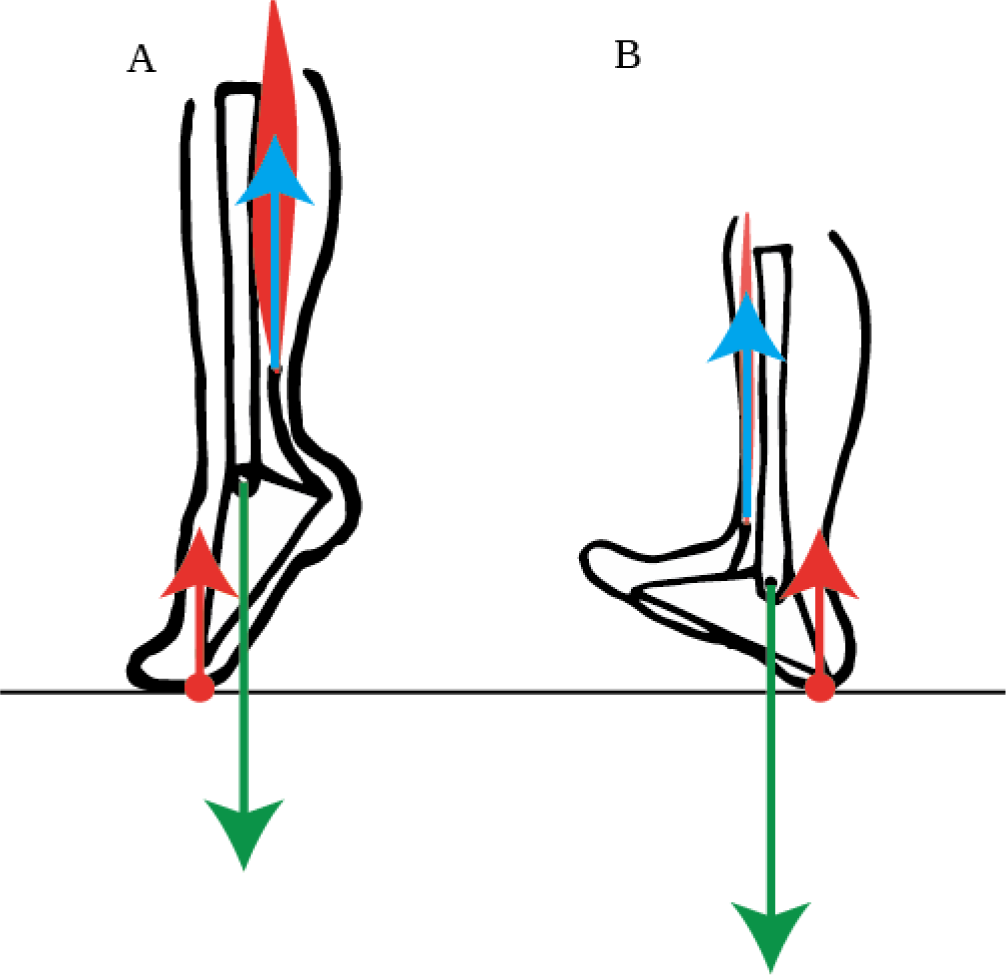
Limited ground reaction torque. A. When the backwards torque exerted by the calf muscles exceeds the product of the person’s weight and the distance between the ankles and toes, the person rises onto their toes. B. When the forwards torque exerted by the shin muscle exceeds the product of the person’s weight and the distance between the ankle and heel, the person rocks onto their heels.

#### 6.1.3 The ground reaction torque is limited by the extent of the foot

However, the CoP cannot move further forwards than the toes. Thus if the contraction of the calf muscle exerts a torque that is larger than the product of the person’s weight and the distance between their ankle and toes, then the foot can no longer remain immobile: the foot must then rotate around the toes. Indeed, when subjects are asked to rise onto their toes, they perform this movement with a burst of contraction of their calf muscles (Fig. 10A, Nardone and Schieppati, 1988). Likewise, shin muscle contraction induces forwards ground reaction torque by shifting the CoP backwards. However, the CoP cannot move further backwards than the heel. Thus when subjects are asked to rock onto their heels, they perform this movement with a burst of contraction of their shin muscle (Fig. 10B, Nardone and Schieppati, 1988).

The potential ground reaction torque is therefore limited by the extent of the foot: the forwards torque is limited to the product of the person’s weight and the distance between the heels and the ankles, and the backwards torque is limited to the product of the person’s weight and the distance between the ankles and toes.

#### 6.1.4 The net torque is limited by the position of the CoM

The ground reaction torque changes instantly when the torques exerted by the lower leg muscles on the foot change, but it is limited by the extent of the foot. The torque of weight on the other hand can only be changed by displacing the CoM forwards or backwards, which cannot be done instantly but first requires the sum of the external forces to accelerate the CoM horizontally. Therefore, at a given instant, the potential net torque that can be induced by muscular contraction is limited by the position of the CoM: the net forwards torque is limited to the product of the weight and the distance between the CoM and the heels, whereas the net backwards torque is limited to the product of the weight and the distance between the CoM and the toes.

### 6.2 Horizontal acceleration of the CoM

We will now consider the horizontal acceleration of the CoM. Since the person’s weight is vertical, only the ground reaction force may accelerate the CoM horizontally.

#### 6.2.1 Acceleration of the CoM induced by muscular contraction

The contraction of the dorsal muscles causes the trunk to rotate backwards around the hips (Fig. 11A). This backwards acceleration of the mass of the trunk implies that the trunk pushes forwards on the hips, which are therefore accelerated forwards. The dorsal trunk muscles do not exert torque on the foot around the ankles, therefore they do not induce a change in the ground reaction force. The person’s rotational momentum around their ankles is therefore unchanged. The increase in backwards rotational momentum around the ankles due to the backwards acceleration of the head must therefore be compensated by an equal increase in forwards rotational momentum due to the forwards acceleration of the hips. Since the head is further from the ankles than the hips are, and since rotational momentum is proportional to distance, this implies that the forwards acceleration of the hips exceeds the backwards acceleration of the head, such that the CoM is accelerated forwards (Fig. 11A).

The contraction of the calf muscles causes the legs to rotate backwards. However, the calf muscles do not exert torque on the trunk around the hips. Therefore, if only the calf muscles contract, then the rotational momentum of the trunk around the initial position of the hips is unchanged: due to its inertia, the trunk therefore rotates forwards in the external frame of reference as the legs rotate backwards. The person therefore flexes at the hips (Fig. 11B). Moreover, the contraction of the calf muscles induces backwards torque from the ground reaction force and therefore increases the person’s backwards rotational momentum around the ankles. The increase in backwards rotational momentum around the ankles due to the backwards acceleration of the hips must therefore exceed the forwards rotational momentum due to the forwards acceleration of the trunk. This implies that the CoM is accelerated backwards (Fig. 11B).

Thus, contracting the dorsal trunk muscles accelerates the CoM forwards (Fig. 11A) and contracting the calf muscles accelerates the CoM backwards (Fig. 11B). In order to accelerate the CoM backwards at the initiation of a movement requiring both calf and dorsal trunk muscle contraction, the calf muscle contraction should therefore precede the dorsal trunk muscle contraction.

**Fig. 11.**
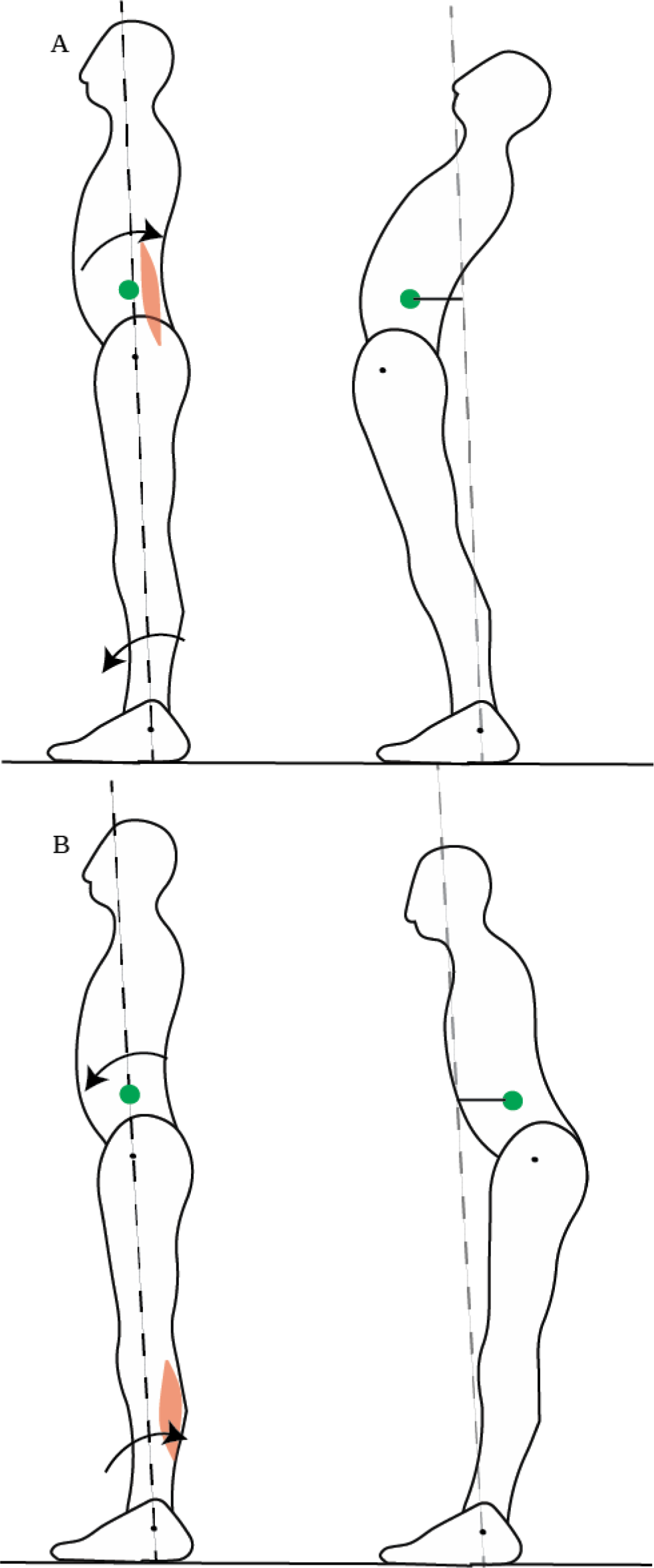
CoM acceleration induced by muscular contraction. A. Dorsal trunk muscle contraction makes the trunk rotate backwards around the hips while the legs rotate forwards around the ankles, accelerating the CoM forwards. B. Calf muscle contraction makes the legs rotate backwards around the ankles while the trunk rotates forward around the hips, accelerating the CoM backwards.

#### 6.2.2 Muscular contractions required for movement

Leaning the trunk forwards requires not only an increase in the trunk’s forwards rotational momentum around the hips, through the contraction of the abdominal muscles, but also an increase in the person’s total forwards rotational momentum around the ankles, through the contraction of the shin muscle. Thus, when leaning forwards, both the abdominals and the shin should be considered as “prime movers”, since they play the same role of providing torque for movement. If we take into account the knee joint, then the same analysis shows that leaning the trunk also requires the contraction of the thigh muscles. Moreover, the initial acceleration of the CoM requires a temporal sequencing of muscular contraction, with the lower leg muscles contracting first, then the thigh and finally the trunk muscles.

In order to straighten up after a platform translation, the person must rotate their entire body backwards around the ankles, keeping their legs and trunk aligned. This movement requires backwards rotational momentum of the body around the ankles, and therefore calf muscle contraction, but also backwards rotational momentum of the trunk around the hips, and therefore contraction of the dorsal trunk muscles. If we take into account the knee joint, then the same analysis shows that straightening up also requires contraction of the dorsal thigh muscles. Moreover, the initial acceleration of the CoM requires a temporal sequencing of muscular contraction, with the lower leg muscles contracting first, then the thigh and finally the trunk muscles.

## 7 Conflict of Interest

*The authors declare that the research was conducted in the absence of any commercial or financial relationships that could be construed as a potential conflict of interest*.

## 8 Author Contributions

Both authors have jointly contributed to the development of the ideas presented and to the writing of the manuscript.

## 9 Funding

This work was supported by Agence Nationale de la Recherche (ANR-14-CE13-0003).

## 10 Acknowledgments

